# INTEGRINS MEDIATE PLACENTAL EXTRACELLULAR VESICLE TRAFFICKING TO LUNG AND LIVER IN VIVO

**DOI:** 10.1101/2020.09.22.309047

**Authors:** Sean L. Nguyen, Soo Hyun Ahn, Jacob W. Greenberg, Benjamin W. Collaer, Dalen W. Agnew, Ripla Arora, Margaret G. Petroff

## Abstract

Membrane-bound extracellular vesicles (EVs) mediate intercellular communication in all organisms, and those produced by placental mammals have become increasingly recognized as significant mediators of fetal-maternal communication. Here, we aimed to identify maternal cells targeted by placental EVs and elucidate the mechanisms by which they traffic to these cells. Exogenously administered pregnancy-associated EVs traffic specifically to the lung; further, placental EVs associate with lung interstitial macrophages and liver Kupffer cells in an integrin-dependent manner. Localization of EV to maternal lungs was confirmed in unmanipulated pregnancy using a transgenic reporter mouse model, which also provided in situ and in vitro evidence that fetally-derived EVs, rarely, may cause genetic alteration of maternal cells. These results provide for the first time direct in vivo evidence for targeting of placental EVs to maternal immune cells, and further, evidence that EVs can alter cellular phenotype.

## INTRODUCTION

Extracellular vesicles (EVs), which can be secreted as exosomes, microvesicles, and apoptotic bodies, are increasingly recognized as a mechanism of intercellular communication in all organisms.^1^ Each type of EV possesses unique cargo and biological activity, which establishes a need for acquiring a basic understanding of their role in physiologic and pathologic processes. Exosomes are 40-150 nm membrane-enclosed EVs that are created by the invagination of the early endosomes to form multivesicular bodies, which then can fuse with either the lysosome or the plasma membrane. When they fuse with the plasma membrane, the intraluminal vesicles are released from the cell as exosomes, carrying bioactive nucleic acids, proteins, and lipids through interstitial space and the circulation to distant cells.^2^ Both the cell of origin and their physiological state have strong influences on exosome quantities and activity.

Among mammalian cells and tissues that shed vesicles, the placenta plays a particularly important role, releasing enormous quantities of EVs in order to communicate with local and distant maternal cells during pregnancy. This is particularly true of hemochorial species, in which the maternal-fetal interface consists of trophoblast cells that juxtapose directly with maternal blood. Humans and mice both possess this type of placentation, with the chorionic villi and labyrinth, respectively, possessing outermost syncytiotrophoblast cells that release EVs into the maternal circulation.^3,4^ Because of this intimate association, placental EVs have the potential to communicate with virtually any tissue in the mother that they are able to access. As gestation advances, placental exosome concentration increases in maternal plasma;^5,6^ additionally, their bioactivity may change in pathological conditions including preeclampsia and gestational diabetes.^7–9^ These changes have spurred interest into possible roles of exosomes in regulating immune and cardiovascular responses to pregnancy and parturition. Further, interest in placental EVs as biomarkers for these and other complications has burgeoned.^10^

In vitro studies have suggested that placental exosomes can be internalized by and affect activity of multiple immune cell types. Placental exosomes can induce migration and cytokine secretion by macrophages,^11^ suppress NK cells and T cells,^12–14^ protect cells against viral infection,^15^ and may promote spiral artery remodeling.^16^ However, only a few studies have queried their trafficking patterns and functional effects in vivo. Tong et al. found localization of human placental exosomes in the lungs, kidney, and liver of mice 24 hours after intravenous injection.^17^ On the other hand, intraperitoneally injected exosomes purified from plasma of pregnant mice traffic to the uterus and cervix, and further, can cross into the fetus.^18^ In this study, we advance our understanding of placental EVs by identifying specific cellular targets of EVs and the molecular mechanisms that influence their trafficking. We also developed a Cre-recombinase reporter system in unmanipulated pregnancy to support these observations. Finally, we provide in vivo and in vitro proof of principle that genetic alteration of maternal cells by placenta-derived exosomes is possible.

## RESULTS

### Pregnant plasma localizes to lung in vivo

Consistent with our prior results,^6^ the concentration of EVs in the plasma of gestation day (GD)14.5 pregnant mice were ~2.4-fold higher than in nonpregnant females (P = 0.015) (Suppl. Fig. 1). As placental EVs are a likely source of this increase, and as they can potentially access all maternal organs via the vasculature, we tested the specificity of their trafficking by comparing localization of plasma EVs from nonpregnant and pregnant mice. To this end, EVs were purified from plasma, labeled with the red fluorescent dye PKH26, and administered into the tail vein of nonpregnant recipient females (Fig. 1A). The lung and liver of recipients were harvested after 30 minutes and evaluated by epifluorescence microscopy. In the lung, only EVs from pregnant mice were readily detectable; EVs from nonpregnant mice were scarce or undetectable (Fig. 1B, C). The same trend did not hold true for EV localization to the liver (Figure 1D, E). These results suggest preferential localization of EVs associated with pregnancy to the lung.

**Figure 1.**
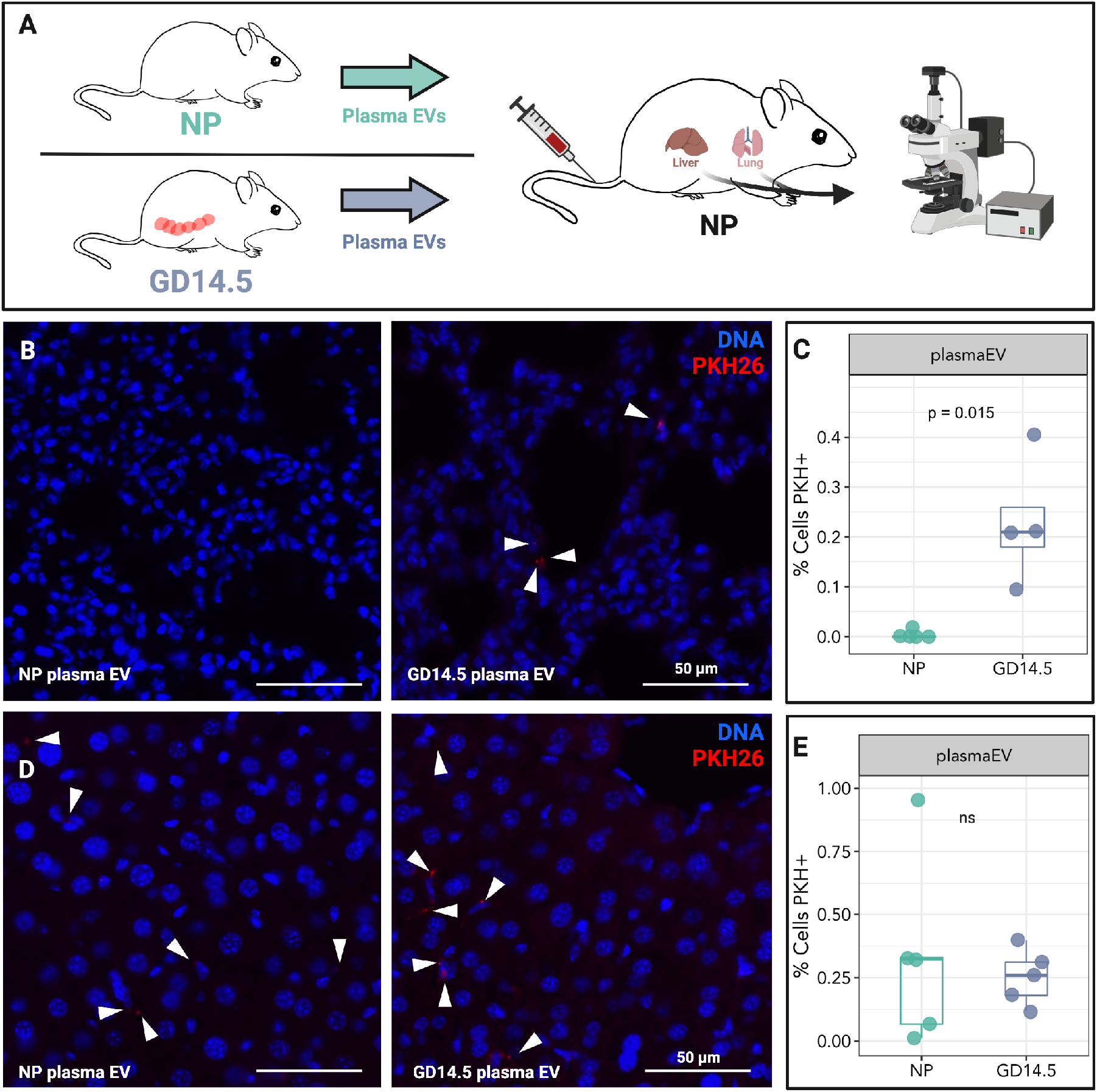
Pregnant plasma EVs traffic to murine lungs and liver in vivo. A. Plasma EVs (2.5×10^10^) isolated from the plasma of nonpregnant (NP) or GD14.5 pregnant mice were labeled with PKH26 and administered i.v. into NP mice. Mice were sacrificed after 30 minutes, and lung and liver were analyzed by epifluorescence microscopy. B. Representative fluorescence microscopy of lung from a mouse treated with plasma EVs from nonpregnant (NP; upper left) or GD14.5 pregnant (upper right) mice (Arrowheads, EVs). C. Quantification of plasma EVs as evidenced by presence of red fluorescence in recipient lung. Dots represent biological replicates, Wilcoxon test. D. Representative fluorescence microscopy of liver from a mouse treated with plasma EVs from nonpregnant or GD14.5 pregnant mice EVs (Arrowheads, plasma EVs). E. Quantification of plasma EVs as evidenced by presence of red fluorescence in recipient liver. Dots represent biological replicates, Wilcoxon test.

### Placental EVs traffic to lung interstitial macrophages in vivo

We next sought to identify the type(s) of cells in the lung with which placental EVs associate. To this end, we developed an explant system in which placental EVs are purified following 18 hours of culture at ambient oxygen concentration (Suppl. Fig. 2). EVs from the supernatant of GD14.5 placental explants were isolated, labeled with PKH26, and administered intravenously into nonpregnant female mice (Fig. 2A).

**Figure 2.**
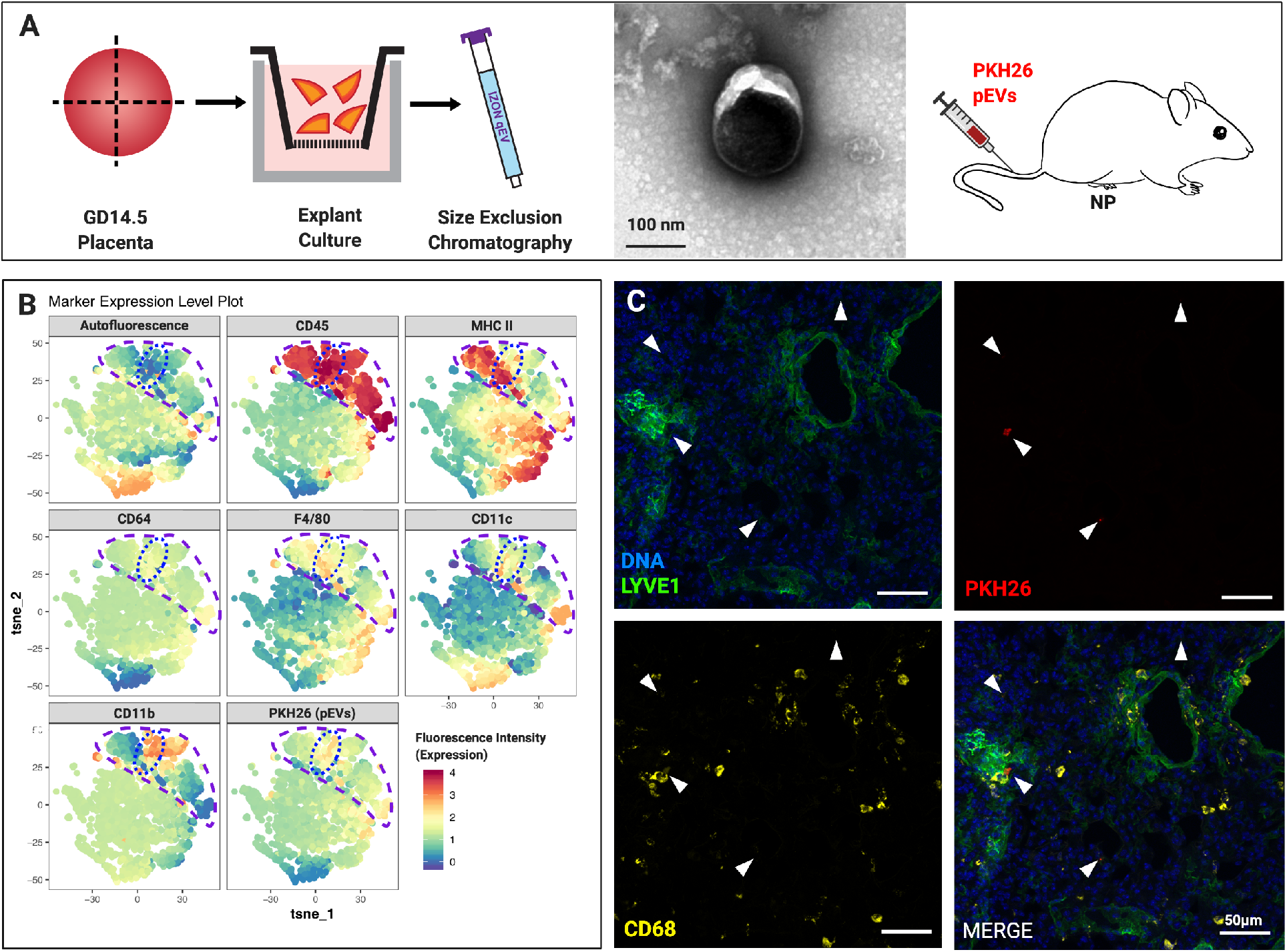
Placental EV trafficking in vivo. A. EVs were purified from the medium of GD14.5 placental explants; a representative TEM is shown. After labeling with PKH26, 2.5×10^10^ EVs were administered i.v. into into nonpregnant females. After 30 minutes mice were sacrificed, and lung and liver harvested and processed for downstream analysis. B. Representative t-SNE analysis of dispersed and murine lung cells after treatment of mice with GD14.5 placental EVs. Cell suspensions were stained with the fluorophore-conjugated antibodies indicated above each panel and analyzed by flow cytometry. C. Immunofluorescence microscopy of lungs from mice treated with GD14.5 placental EVs. Arrows, pEV foci (red), one of which in this image colocalizes with LYVE1 (green) and CD68 (yellow). Representative images of random fields chosen from three independent experiments.

After 30 minutes, lungs were harvested and dispersed for flow cytometry. For unbiased identification of immune cells in the lung that associated with PHK26, we used t-distributed stochastic neighbor embedding (t-SNE). This approach identified CD45+ cells, and within these, cells positive for PKH26 (Fig. 2B). PKH26-positive cells were also CD45+, MHCII^hi/lo^, F4/80+, autofluorescence (AF)-, CD11c^lo^, and CD64+ – all characteristic of interstitial macrophages (Fig. 2B).^19^ We confirmed this result first by immunofluorescence microscopy, in which PKH26 colocalized with LYVE1+ and CD68+ interstitial macrophages (Fig. 2C).^19^ Second, using forward-gating strategy to complement t-SNE analysis,^20^ we found that PKH26+ placental EVs localize predominantly with interstitial, but not alveolar, macrophages (Fig. 3). Confocal microscopy of liver tissue from the same animals revealed localization of placental EVs within CD31+ endothelial cells and F4/80+ Kupffer cells (Suppl. Fig. 3A, 3B).

**Figure 3.**
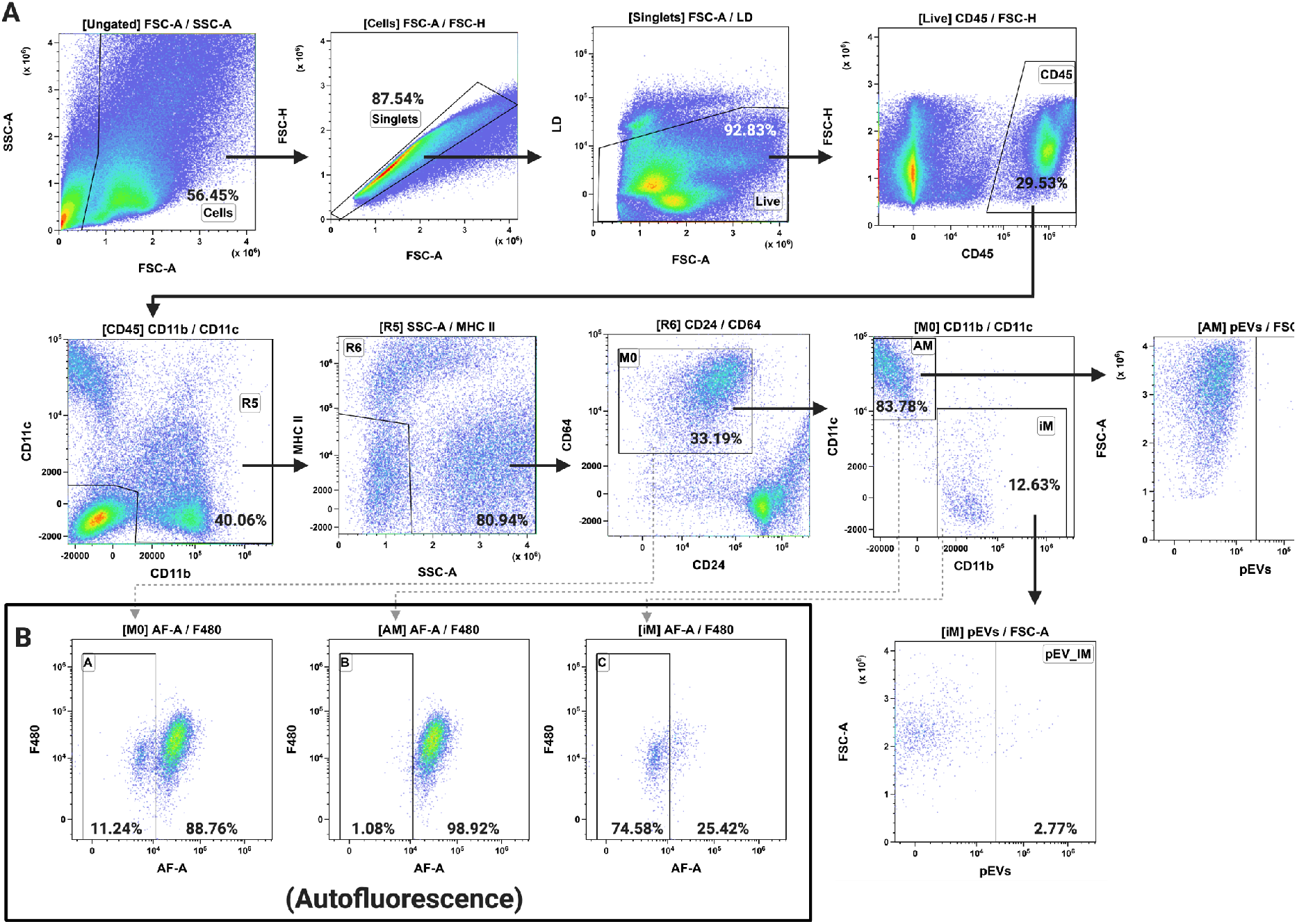
Flow cytometric analysis of pEV trafficking in murine lung. A. Representative flow cytometry plots identifying macrophages (M0), alveolar macrophages (AM), and interstitial macrophages (iM) from murine lung after treatment of mice with GD14.5 placental EVs. B. Autofluorescence signature of macrophage, alveolar macrophage, and interstitial macrophage populations.

### Outer membrane proteins including integrins influence trafficking of placental EVs to the lung and liver

We next asked whether outer membrane components, specifically integrins, influence trafficking of placental EVs, as these proteins can mediate the trafficking of tumor-derived EVs to specific organs.^21^ A survey of multiple integrins revealed that placental EVs express ITG *α*3, *α*V, *α*5, *β*1, and *β*3, whereas ITG *β*6 was not detected in EVs despite its expression in the placenta (Fig. 4A). As expected, proteinase K treatment of EVs abolished immunoreactivity for all membrane-associated integrins examined, although neither expression of the exosome marker CD9 nor exosome morphology were affected (Fig 4A, B).

**Figure 4.**
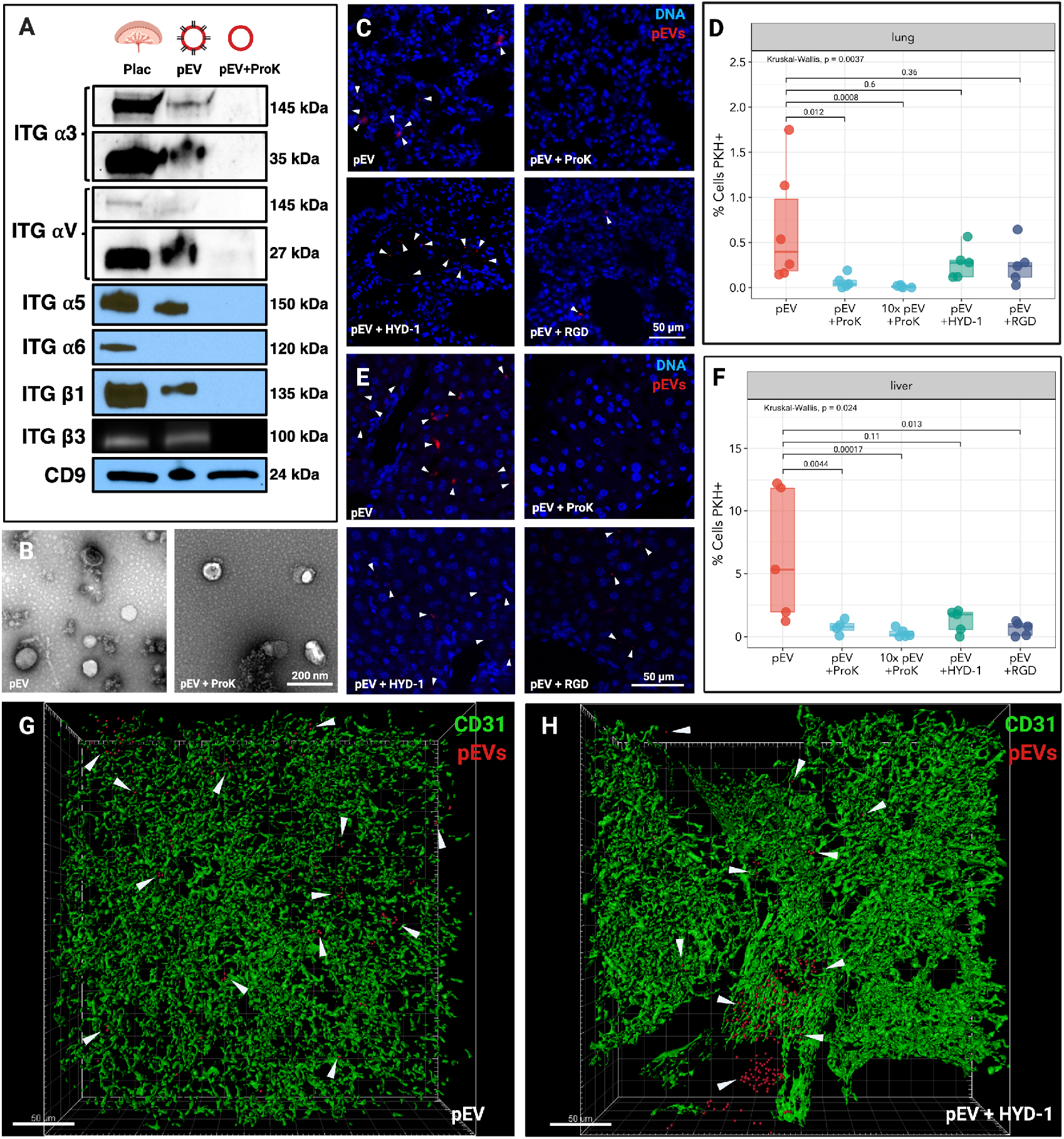
Integrins mediate placental EV localization to murine lung and liver. A. Western Blot analysis of integrin expression in murine placenta, placental EVs (pEV), and placental EVs treated with proteinase K (pEV + proK). B. Representative transmission electron microscopy of untreated EVs (left) or EVs treated with proteinase K (right). C. Confocal microscopy of murine lung after treatment of mice with GD14.5 placental EVs (upper left), proteinase K-treated EVs (upper right), or EVs pre-incubated with HYD-1 (lower left) or RGD (lower right) peptides. D. Quantification of EV positive cells in lungs treated with placental EVs. E. Confocal microscopy images of the liver after treatment of mice with placental EVs (upper left), proteinase K-treated EVs (upper right), or EVs pre-incubated with HYD-1 (lower left) or RGD (lower right) peptides. F. Quantification of EV-positive cells in liver of mice treated with placental EVs. G. CUBIC-cleared lung from mouse treated with placental EVs. White arrowheads represent placental EV foci. H. CUBIC-cleared lung from mouse treated with HYD-1-pretreated EVs.

To determine whether pEV localization to lung and liver was mediated by outer membrane proteins, EVs were digested with proteinase K, labeled, and administered into nonpregnant mice. Proteinase K abolished localization of EVs to the lung and liver (Fig. 4C-F) of EV-treated mice. To account for the possibility that proteinase K caused degradation of EVs, we also administered 10-fold more EVs, and still observed vastly reduced numbers of EVs in liver and lung (Fig. 4D, F). Consistent with this observation, treatment with proteinase K to remove outer membrane proteins reduced the localization of EVs to lung interstitial macrophages (Suppl. Fig. 4).

To test the hypothesis that integrins mediate placental EV targeting to the lung, we pre-incubated EVs with RGD and HYD-1 peptides, which block binding of integrins *α*5*β*1/*α*V*β*3 and *α*3*β*1 respectively, to their substrates^22,23^ prior to intravenous administration into nonpregnant recipients. Mice were sacrificed 30 minutes after treatment, and lung and liver were analyzed by fluorescence microscopy (Fig. 4 C, E). Although proteinase K ablated localization of placental EVs to both lung (Fig. 4C, upper right panel) and liver (Fig. 4E, upper right panel), placental EVs remained detectable in both tissues after pretreatment with integrin-blocking peptides (Fig. 4C, E, lower left panels). In the lung, neither peptide significantly reduced the total numbers of placental EVs (Fig. 4D). In the liver, however, RGD pretreatment resulted in a significant reduction of placental EV localization (Fig. 4F).

Although the total numbers of placental EVs that localized to the lung were not reduced by HYD-1, we noticed that the tissue distribution of the vesicles was altered, with EVs appearing to remain within large vessels (Fig. 4C, lower left panel). To better characterize the effect of HYD-1 treatment on placental EV localization to the lung, 200-micron lung sections were cleared by CUBIC and imaged by 3D confocal microscopy.^24^ Strikingly, we saw localization of placental EVs exclusively within the vasculature, suggesting that they were blocked from entering the interstitial tissue (Fig. 4G, H).

Since HYD-1 pretreatment inhibited rapid migration of placental EVs to lung interstitium, we asked whether this inhibition is sustained. Using Li-Cor whole organ imaging as a high throughput screening method, we labeled EVs with near-infrared (NiR) dye, administered them i.v. into nonpregnant recipients, and quantified vesicle localization (Fig. 5A). Untreated EVs remained detectable in the lung after 24 hours, but pretreatment with HYD-1 significantly decreased their presence (Fig. 5B). RGD pretreatment tended to reduce localization to the lung at 24 hours, but the result was not statistically significant.

**Figure 5.**
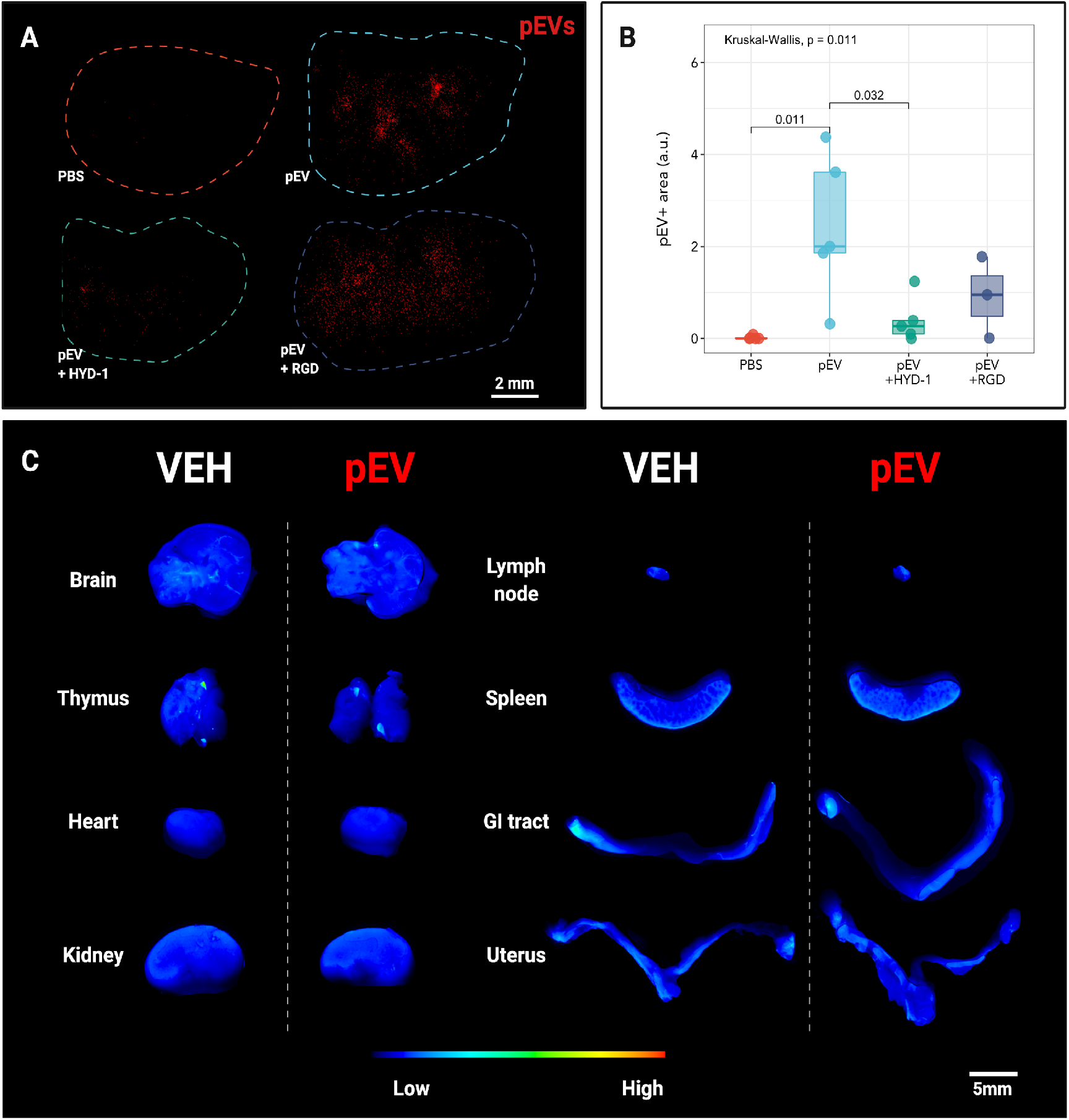
Integrin *α*3*β*1 mediates placental EV trafficking to the lung. A. Near-infrared imaging of whole lung after in vivo treatment with labeled placental EVs after 24 hours. Image is representative of three independent experiments. B. Quantification of NIR placental EV positive area from whole lung treated with placental EVs. a.u., arbitrary units; each point represents one biological replicate. C. Representative whole organ NIR imaging of mice treated with control medium (VEH) or placental EVs (pEV) after 24 hours. Representative of three independent experiments.

We also imaged other organs including, the brain, thymus, heart, kidney, paraaortic lymph node, spleen, small intestine, uterus, and ovaries, again reasoning that EVs in the maternal vasculature can potentially access any maternal organ. Twenty-four hours after administration of NIR-labeled placental EVs, we did not observe any significant changes in NIR signal relative to the vehicle control treatments (Fig. 5C). This result suggests that placental EVs are not detectable in these tissues using this model.

### Placental EVs localize to maternal lung in unmanipulated pregnancy and may genetically modify maternal lung cells

The above studies support the notion that placental EVs traffic to the maternal lung. However, a single bolus of purified EVs does not recapitulate the continual release of EVs from the placenta nor their sustained presence in maternal circulation. To address this limitation, we generated a model in which fetal EVs could be detected within the context of normal pregnancy. mTmG mice^25^ express a fluorescent reporter construct that, in the absence of Cre recombinase, allows constitutive expression of membrane-targeted red fluorescent protein, tandem dimer Tomato (mTomato). In the presence of Cre, the mTomato locus is excised, resulting in the expression of membrane-targeted enhanced green fluorescent protein (EGFP; mGFP). By mating mTmG females with CMV-Cre (Fig. 6A), female fetuses inherit both the CMV-Cre and mTmG loci, and thus express mGFP in all tissues, including the placenta. Male embryos inherit the wild type (WT) locus and continue to express the mTomato red fluorescent protein. Confocal microscopy (Fig. 6B) and western blot analysis (Fig. 6C) confirmed expression of mGFP in both placentas and placental EVs from female CMV-Cre;mTmG but not WT;mTmG fetuses. Both CMV-Cre;mT/mG and WT;mT/mG placentas and pEV samples expressed the control protein, TSG101.

**Figure 6.**
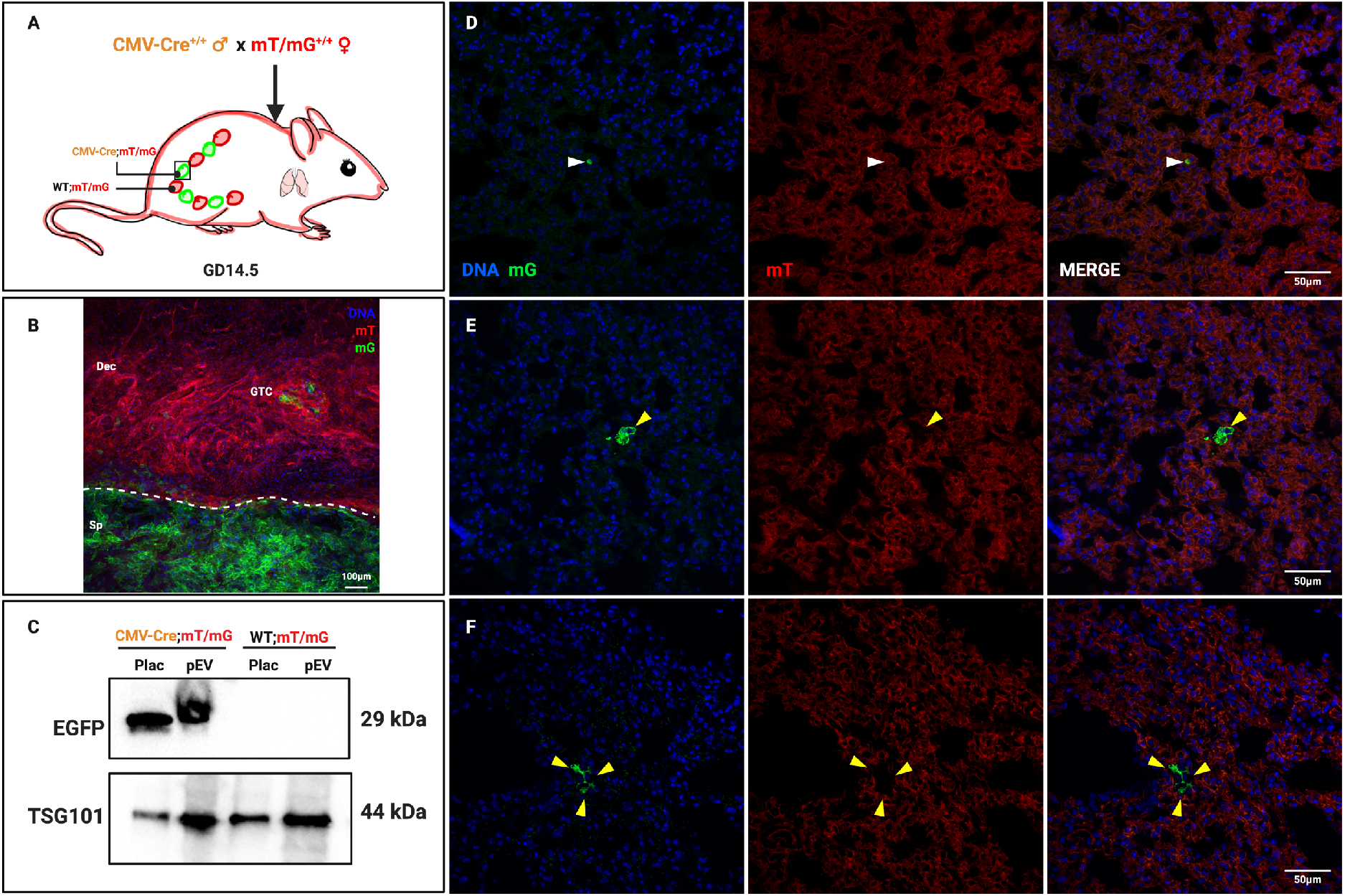
In Vivo Localization of mGFP in CMV-Cre Mated Maternal mT/mG Lung. A. mT/mG females mated to a CMV-Cre males give rise to female pups ubiquitously expressing mGFP and male pups expressing mTomato. B. GD14.5 utero-placental interface of a female fetus from a mT/mG x CMV-Cre mating. Fetal mGFP-expressing placental cells (Sp; spongiotrophoblast) are readily distinguishable from maternal mTomato-expressing decidual cells (Dec); invading glycogen trophoblast (GTC) can also be seen. The dashed line demarcates the fetomaternal interface. Representative image of three independent experiments. C. Western blot of GD14.5 placenta (Plac) and placental EVs (pEVs) from mT/mG females mated with CMV-Cre or WT males. D. Confocal image of punctate mGFP-positive foci localization in maternal mT/mG GD14.5 lung after mating to a CMV-Cre male. Representative of n=5. E. **E, F.** Confocal localization of mG+ recombined cells in maternal mT/mG GD14.5 lung after mating to a CMV-Cre male. Images are representative of five biological replicates. White arrows; punctate mGFP-positive foci. Yellow arrows; mGFP-positive, recombined cells.

Using this model, we asked whether fetal/placental mGFP+ EVs are carried to the lung. Using confocal microscopy, we readily detected mGFP fluorescence in maternal lung from GD14.5 CMV-Cre-mated mTmG females (Fig. 6D). Most of the signal we observed was punctate and associated with maternal mTomato-expressing cells, similar to what was observed when labeled placental EVs were administered exogenously. Interestingly, we also observed mGFP expression that was clearly membrane-associated, surrounding distinct DAPI-stained nuclei (Fig. 6E, F). These cells did not express mTomato, indicating that they had undergone Cre-mediated recombination. We observed mGFP-associated foci and cells only in mT/mG dams mated to CMV-Cre males, and not in those mated to WT males, therefore ruling out random recombination of the mT/mG locus in our experimental animals.

### Placental EVs can induce Cre-mediated recombination in vitro

Recombined mGFP-positive cells observed in the lungs could represent fetal cells that trafficked from the fetus/placenta to the maternal lungs (fetal microchimerism).^26–28^ Alternatively, they could be maternal cells that underwent Cre-mediated recombination as a result of EVs carrying Cre mRNA or protein to recipient cells.^29,30^ To test the latter possibility, we cocultured bone marrow-derived dendritic cells (BMDC) from mTmG mice with GD14.5 placental explants from CMV-Cre (colorless) or WT mice (Fig. 7A).

**Figure 7.**
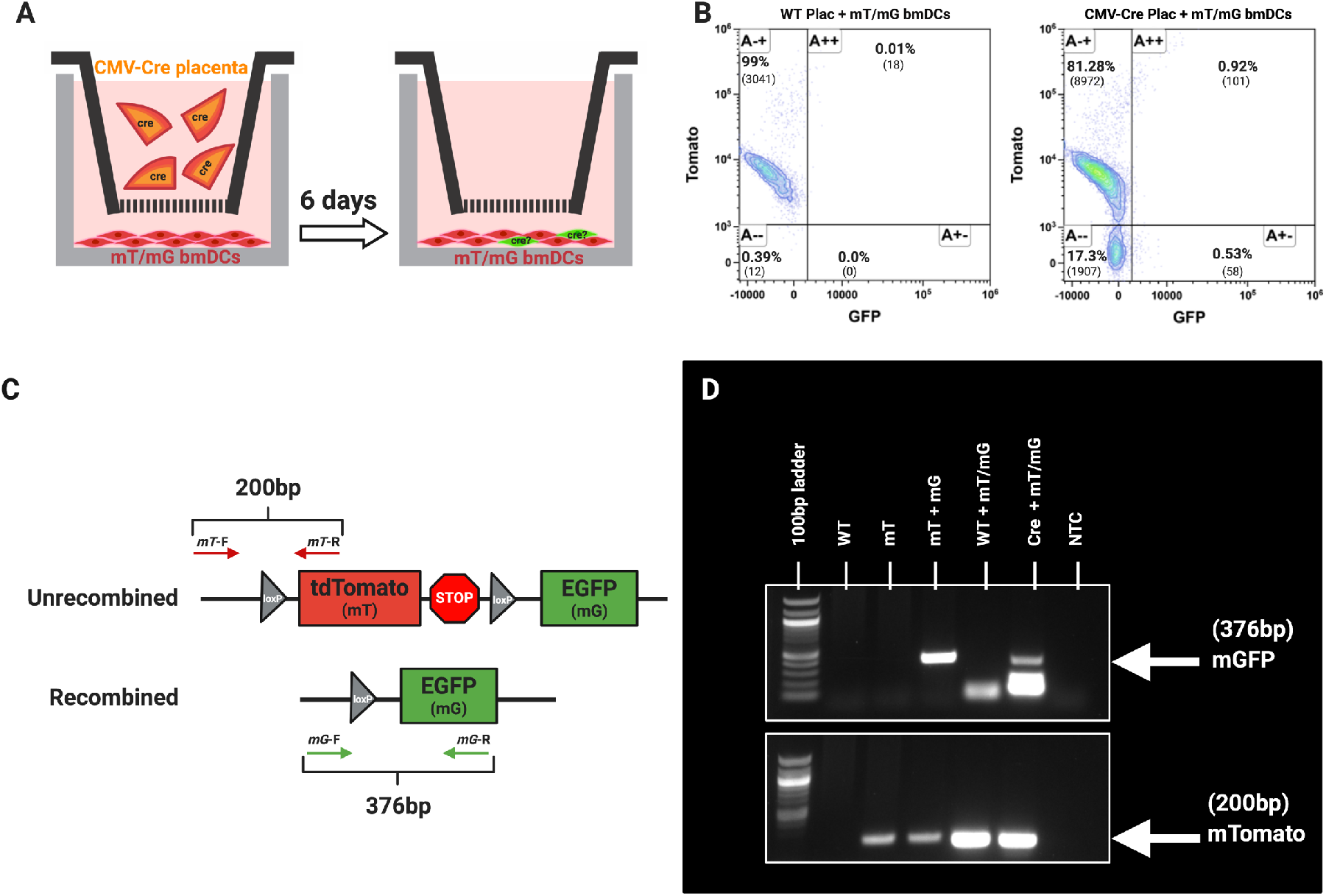
In Vitro Transfer of Placental Cre to Recipient Reporter Cells. A. mT/mG BMDCs were cultured with (colorless) GD14.5 CMV-Cre placentas for 24 hrs and cultured for an additional five days. B. Representative flow cytometry plots of mT/mG BMDCs cocultured with WT (left) or CMV-Cre (right) placentas. C. Schematic diagram of forward (mT-F/mG-F) and reverse (mT-R/mG-R) primers for identification of unrecombined mTomato and recombined mGFP. PCR amplification of genomic DNA with these primers results in PCR products of 376 bp for non-recombined cells and 200 bp for recombined cells. D. Genotyping electrophoresis gel of mGFP (top) and mTomato (bottom) loci from genomic DNA. WT: tail DNA from a wild type mouse; mT: tail DNA from an mTmG mouse in the absence of Cre; mT+mG: tail DNA from a fully mTmG mouse; WT plac+mT/mG: DNA from dendritic cells cocultured with WT placenta; CMV-Cre plac + mT/mG BMDC: DNA from dendritic cells cocultured with CMV-Cre placenta; NTC, no template control. Representative gel image of five independent experiments.

Explants were separated from the dendritic cells by a 70 μm insert such that EVs released into the medium could access the underlying BMDC. After 18 hours, placentas were removed, and BMDC were cultured for an additional five days to allow sufficient time for Cre recombination to occur.^25,30^ The BMDC were screened for recombined mGFP positive cells by flow cytometry. While no changes in mTomato expression occurred in BMDC cultured with WT placentas (Fig. 7B, left), culture with CMV-Cre placentas resulted in a downward shift of events into a new mTomato-negative/mGFP negative population and an increase in the proportion of mTomato-negative/mGFP-positive population (Fig. 7B, right).

To confirm these results, we isolated the DNA from the cocultured BMDC and performed PCR using primers designed to differentially amplify the non-recombined and recombined mTmG locus (Fig. 7C).^31^ We observed the mGFP amplicons in DNA isolated from BMDC cocultured with CMV-Cre, but not WT placentas (Fig. 7D). Finally, confocal microscopy conducted in parallel revealed the presence of cell-associated and non-cell-associated mGFP in BMDC co-cultured with CMV-Cre placentas, but not WT placentas (Suppl. Fig. 5), which we confirmed using immunofluorescence with an anti-GFP antibody (Suppl. Fig. 5B, D). Interestingly, the morphology of dendritic cells co-cultured with CMV-Cre placentas revealed large clusters of cells that were not observed when the cells were co-cultured withWT placentas (Suppl. Fig. 5D). Our results demonstrate a proof of concept model for identifying recipient cells that internalize placenta EVs and provides an additional mechanism for studying how placental EVs induce biological effects on recipient cells.

## DISCUSSION

Since the discovery of EVs and their effects on distant cells, research interest in the role of placental EVs during pregnancy has greatly increased. The total quantity and concentration of EVs in maternal plasma rise across gestation, with the placenta contributing significantly to this increase.^5,6^ While a number of effects of placental EVs have been suggested, few studies have attempted to quantify their biodistribution in vivo. In this study, we show that placental EVs traffic to the lung and the liver. Further, that trafficking to the lung appears to be specific to placental EVs and pregnancy, as only plasma EVs isolated from pregnant dams, but not those from non-pregnant mice, localized to the lung.

Our results align with those of Tong et al., who found that human placental EVs administered into mice also localize to the lung and liver.^17^ In our study, we add to these findings by using homologous adoptive transfer of murine EVs to identify the cellular targets of placental EVs in these tissues. Using multiparameter flow cytometry together with both targeted and untargeted analyses, as well as immunofluorescence microscopy, we found that murine placental EVs target lung interstitial macrophages and liver Kupffer cells. While this result is not especially surprising, it reveals for the first time the bona fide in vivo targets of EVs and aligns with earlier data showing that human trophoblast EVs are internalized by macrophages in vitro.^11^

EVs derived from Swan-71 cells, an extravillous trophoblast cell line, were previously shown to induce migration and proinflammatory cytokine production by cultured macrophages.^11^ Similarly, Southcombe et al. showed that syncytiotrophoblast microparticles, possibly including exosomes, induce pro-inflammatory cytokine release from monocytes.^32^ While the precise role of this induction of cytokines by monocytes is uncertain, studies have supported the notion that pregnancy is associated with a shift in inflammatory environment in general, with a bias towards a proinflammatory milieu in early and late gestation, and anti-inflammatory milieu during mid-gestation.^33^

Whether these observations hold true for the effects of EVs on interstitial macrophages of the lung, which arise from monocytes and play a role in lung homeostasis,^19,34^ is as of yet not known, and the impact of placental EVs on the pulmonary physiology and pathology of the lung during pregnancy warrants investigation. Pregnancy alters maternal pulmonary function dramatically, with up to 20% increase in maternal oxygen consumption by term,^35^ and differential susceptibility to respiratory pathology caused by influenza virus;^36,37^ varicella virus;^38,39^ asthma;^40^ and cigarette smoking.^41^ The notion that placenta- and pregnancy-associated EVs mediate these physiological and pathological adaptations to pregnancy via effects on pulmonary immune cells including macrophages is untested but intriguing.

Another important function of macrophages is antigen presentation. Maternal T cells are made aware of fetal antigen through indirect antigen presentation by maternal antigen presenting cells,^42^ which may include macrophages. Maternal T cells that recognize fetal antigen do not mount an adverse immune response to fetal antigen, even when artificially stimulated with high concentrations of adjuvant,^42^ suggesting that antigen presenting cells in pregnancy convey powerful tolerogenic signals. We and others have postulated that EVs are the source of these tolerogenic signals, possibly through cargo that include potent suppressors such as PDL1 and FASL, as well as fetal antigen itself.^13,14,43^ Surprisingly, our current results show minimal or no localization of placental EVs to the maternal spleen and lymph nodes, where fetal antigen recognition by maternal T cells occurs,^42,44^ but rather to lung interstitial macrophages and Kupffer cells of the liver. Lung interstitial macrophages together with resident dendritic cells are potent antigen presenting cells.^19^ Future work can examine the role of placental EVs in antigen presentation by these macrophages as a possible mechanism for maternal immune system exposure and tolerance to fetal-placental antigens.

Our results highlight possible roles for integrins on placental EVs. The integrin profile in placental EVs did not fully recapitulate integrin expression in the placenta: integrins *α*3, *α*V, *α*5, *β*1, and *β*3 were expressed in both placenta and EVs, while α6 was found only in the placenta. This suggests that proteins are selectively loaded into placental EVs during their biogenesis, a notion supported by numerous other studies.^45^ Additionally, removal of surface proteins, including integrins, using proteinase K pre-treatment disrupted the trafficking pattern of placental EVs. Further, pre-incubation of EVs with RGD peptide, which blocks ITG *α*5*β*1 and *α*V*β*3 binding to its receptor, fibronectin, abrogated appearance of placental EVs in the liver, which is rich in fibronectin.^46^ Similarly, HYD-1, which blocks ITG *α*3*β*1 binding to the pulmonary basement membrane glycoprotein laminin (LN)-5, ^23,47^ prevented entry of placental EVs into the lung. This observation was supported by 3-dimensional imaging, which highlighted association of EVs with lung endothelial cells, and confirmed that EVs remain restricted to large vessels in the lung when pre-incubated with the HYD-1 peptide.

Placental EV adhesion and extracellular matrix proteins may also play an important role in metastasis of choriocarcinoma. Trophoblast cells – the source of EVs in our studies – share many properties with cancer cells, including epithelial-to-mesenchymal transition and invasion into adjacent tissues during the physiological process of embryo implantation.^48^ When trophoblast cells become malignant, the lung is a primary site for metastasis. In a murine xenograft model of cancer, EVs derived from metastatic breast cancer tumors selectively trafficked to the lung and liver, which served to establish a niche for future metastasis to these tissues.^21^ Moreover, integrins played a major role in the selective targeting of EVs to the lung.^21^ Thus, placental EV trafficking to the lung could play an unfortunate role in establishing a niche, parallel to that found for other metastatic tumors.

A limitation of our studies is that bolus injection of purified EVs does not recapitulate the physiological process of continuous EV release during pregnancy. We sought to address this caveat in multiple ways. First, we used a quantity of EV that mimicked quantities found during pregnancy.^6^ Second, we administered EVs into the tail vein, reasoning that this route most closely mimics hematogenous release of placental EV *in situ*, as both this and the uterine vein ultimately drain into the inferior vena cava. Thus, although we were surprised that our results did not show trafficking of placental EVs to the uterus, we believe that intravenous administration simulates the hematogenous route that EVs travel during pregnancy.

A third way we addressed the limitations of bolus EV administration was to develop an in vivo model without the need to isolate, label and administer them. In this system, fetal tissues and EVs derived thereof express a mGFP reporter that enables their in vivo tissue and cellular targets. EVs could be detected as punctate foci within the maternal lung pregnancy, mimicking the pattern observed after intravenous injection. We propose that this model is most representative date of continuous placental EV release and trafficking, and that it will allow further studies on the effects of EVs on maternal physiology in vivo.

We also identified whole cells in the lung that had recombined to express mGFP but not mTomato. Recombined mGFP-expressing cells in the maternal lung may arise from either or both of two mechanisms: genetic recombination of maternal lung cells by placental EVs, or fetal microchimerism. In our model, Cre recombinase is inherited paternally by the fetus, causing recombination of the mTmG locus that switches cells from red to green fluorescence. Using similar models, others have shown that Cre mRNA is carried by exosomes secreted by cancer cells, and that these exosomes could thus induce recombination of neighboring or distant cells.^30^ To test whether this is possible in principle, we used a model system in which red fluorescent mTmG dendritic cells were co-cultured with CMV-Cre placentas. Data generated by immunofluorescence microscopy, flow cytometry, and genetic analysis support the notion that shed vesicles from the explants induced recombination in the dendritic cells.

A second possible explanation for our observation of recombined cells in the mother is fetal microchimerism. Fetal cells that traffic into maternal tissues was first described with the presence of fetal trophoblast cells residing in the maternal lung over 125 years ago^49^ and has been confirmed by numerous studies since.^26,50,51^ Fetal microchimerism is persistent: male fetal progenitor cells are detectable in maternal blood for decades postpartum.^50^ Murine models suggest that fetal nucleated cells are detectable in maternal lung;^26,27^ the tissue specificity of microchimerism may depend on maternal physiological and pathological conditions.^52^ To elucidate whether genetic recombination of maternal cells and/or fetal microchimerism is responsible for the appearance of recombined cells in our model, ongoing studies in our lab seek to identify Cre recombinase protein or mRNA in EVs and will use in vivo approaches to analyze directly whether placental EVs can induce genetic changes in maternal cells.

Collectively, we have established that placental EVs preferentially traffic to maternal pulmonary interstitial macrophages and Kupffer cells in vivo. We demonstrate that integrin *α*3*β*1 is necessary for localization to the lung interstitial tissue, and that integrins *α*5*β*1 and *α*V*β*5 are necessary for localization to the liver. Future work will seek to identify how placental EVs influence maternal interstitial macrophages and Kupffer cell function, as well as the physiology of the lung and liver, during pregnancy. Additionally, we demonstrate a new model for further expanding the field of understanding placental EV interactions in vivo and provide a framework for visualizing maternal-fetal interactions without the use of exogenous purification and labeling, providing a strong advantage to traditional methods of tracking placental EV kinetics in vivo in a manner consistent with their physiological release across pregnancy.

## METHODS

### Animal experiments

All animal experiments were approved by the Michigan State University Institutional Animal Care and Use Committee (protocol no. 201800176). Wildtype C57BL/6J, B6.129(Cg)-Gt(ROSA)26Sor^tm4(ACTB-tdTomato-EGFP)Luo^/J (mT/mG), and B6.C-Tg(CMV-Cre)1Cgn/J (CMV-Cre) mice were purchased from the Jackson Laboratory (Bar Harbor, ME) and housed in temperature-controlled, 12:12 hr light/dark cycle rooms with food and water available ad libitum. Mice used in experiments were aged 6-12 weeks and timed matings were performed, in which the presence of a vaginal plug was designated gestational day (GD) 0.5. For plasma isolation mice were anesthetized with 3% isoflurane infused with oxygen (2 L/min), and. whole blood was collected by cardiac puncture into EDTA-containing tubes. For injection of EVs and subsequent tissue harvesting, mice were anesthetized and EVs administered via the caudal vein, and tissues were harvested after 30 minutes or 24 hours. All mice were euthanized under anesthesia by cervical dislocation and bilateral pneumothorax.

### Materials and Reagents

Chemicals, reagents, and kits were obtained from Thermo-Fisher (Rockford, IL) or Sigma-Aldrich (St. Louis, MO) unless otherwise stated. Filters and membranes were obtained from Millipore (Burlington, MA), and tissue culture materials were purchased from Corning (Corning, NY). Antibodies and other key reagents are listed in the key resources table. All figures were created using www.BioRender.com.

### Placental Explant Culture

EV-free media (RPMI 1640, 50 *μ*M *β*-mercaptoethanol, 100 *μ*g/ml Penicillin/Streptomycin, 1mM Sodium pyruvate, 10% fetal bovine serum) was ultracentrifuged at 100,000 x g for 18 hrs at 4°C and sterilized through a 0.22 *μ*m filter. Placental explant culture was performed as previously described;^53^ briefly, individual GD14.5-16.5 placentas were cut into four pieces and placed in 74 *μ*m mesh 15 mm net well inserts in a 12-well plate filled with 3 ml of EV-depleted media for 18 hours at 37°C, 5% CO2. To determine optimal culture conditions, viability of tissue explants was assessed by a board-certified veterinary pathologist (DA) by examination of hematoxylin and eosin stained sections using a double blinded scoring system for placental tissue necrosis, using a scale of 0 (no necrosis) to 5 (complete necrosis). Following culture, the culture supernatant was centrifuged twice at 500 xg and 2000 xg for 15 minutes to pellet cells and cellular debris, respectively. Supernatant was filtered through a 0.22*μ*m PVDF membrane filter and concentrated to a volume of 500*μ*l at 3320 xg using 100 kDa molecular weight cut off Vivaspin centrifugal ultrafiltration columns (Sartorius, Stonehouse, UK). The resulting filtrate was stored at −80°C until processed for EV isolation.

To ensure EVs were isolated from high quality tissue, we first tested the viability of the cultured explants under various conditions, as well as the quality and quantity of EVs (Suppl. Fig. 2A). We saw no significant differences in necrosis or EVs between freshly isolated and cultured placentas, although the latter exhibited more variation. We observed the highest yield of EVs from placentas cultured for 24-hour at ambient oxygen, and therefore used these conditions for further studies. For low oxygen culture, tissues were placed in a sealed gas chamber and purged of ambient air and charged with a gas mixture containing 8% oxygen, 5% carbon dioxide before being sealed and placed in an incubator.

### Extracellular vesicle isolation and labeling

Extracellular vesicles were isolated from plasma and concentrated supernatant of explant cultures using qEVsingle or qEVoriginal size exclusion chromatography columns (Izon Sciences, Medford, MA) following the manufacturer’s instructions. Briefly, plasma or supernatant were loaded onto columns equilibrated with PBS and fractions 6-8 or 7-9 (200*μ*l for qEVsingle, 500*μ*l for qEVoriginal columns, respectively) were collected and concentrated to 50*μ*l using Amicon Ultra 4 10kDa centrifugal filters for further analysis.

Purified EVs were labeled with PKH26 lipophilic red fluorescent dye (Sigma-Aldrich, St. Louis, MO) using the manufacturer’s instructions. Briefly, purified EVs or PBS (negative control for injection) were diluted in Diluent C and pipetted into ultracentrifuge tubes containing PKH26 dye and Diluent C. The labeling reaction was stopped after 5 minutes with the addition of 10% BSA and EV-depleted culture medium. To eliminate excess unbound dye, 0.971 M sucrose was pipetted beneath the labeled EVs and centrifuged for 1.5 h at 150,000 xg at 4°C as described.^54^ The labeled EVs were washed and concentrated with Amicon Ultra 4 10kDa centrifugal filters as described above. Prior to administration, we separated any remaining free dye from labeled EV by centrifugation over a sucrose gradient; as a negative control, PKH26 dye alone was processed and administered in parallel.

For experiments in which whole organ imaging was performed, purified EVs were labeled with 0.246mM (DiIC18(7) (1,1’-Dioctadecyl-3,3,3’,3’-Tetramethylindotricarbocyanine Iodide)) DiR for 3 minutes and ultracentrifuged on a sucrose cushion as described above. After centrifugation, the labeled vesicles were loaded on IZON single columns and fractions 6-8 were collected and concentrated with 10kDa concentrators as described above and used for downstream analysis.

### Nanoparticle tracking analysis

EVs were quantified using NanoSight NS300 (Malvern Panalytical, Westborough, MA) as described previously with modifications.^6^ Briefly, concentrated samples were diluted 1:500-2000 in PBS and loaded into a 1 ml syringe at a flow rate of 50. For each sample, five 20-second videos were recorded at camera level 12 and a detection threshold of 4 before data raw data were analyzed with the tidyNano R package.^6^

### Transmission electron microscopy

For transmission electron microscopy, concentrated EVs were fixed with 4% paraformaldehyde in PBS and placed on formvar-carbon coated grids and counterstained with 2.5% glutaraldehyde and 0.1% uranyl acetate in PBS. Negative TEM was performed on a JEOL100CXII (JEOL, Peabody, MA).

### Proteinase K and inhibitory peptide treatment

For proteinase K treatment, placental EVs were treated with 10*μ*g/ml proteinase K in PBS; for controls, proteinase K was added to PBS alone. Samples and controls were incubated for 10 minutes at 37°C before heat inactivation at 65°C for 10 minutes followed by addition of 100*μ*M phenylmethylsulfonyl fluoride (PMSF) protease inhibitor.

For inhibitory peptide experiments, RGD peptide was purchased from Sigma-Aldrich and HYD-1 (KIKMVISWKG) was synthesized (Genscript, Piscataway, NJ). All reagents were 95% or greater purity and resuspended in deionized water, aliquoted, and frozen. Labeled pEVs were incubated with 0.6 *μ*M of inhibitory peptide and incubated at 37°C for 30 minutes before intravenous administration into recipient mice.

### mT/mG BMDC explant co-culture

Bone marrow derived dendritic cells (BMDC) were prepared from mT/mG mice as described with modifications.^55^ Ten million bone marrow cells/well were plated overnight in medium containing 20ng/ml murine GM-CSF(BioLegend, San Diego, CA), replenishing the medium daily for three days, then culturing for an additional three days before downstream analysis.

For co-culture with placental explants, 100,000 BMDC were seeded in 12-well plates, and CMV-Cre or WT placentas were added on Netwell inserts as described above. Following overnight culture, the placentas were removed, and the BMDC were cultured for an additional five days to allow for sufficient time for recombination,^25^ replenishing the media every other day. For microscopy, BMDC were seeded onto coverslips before placental co-culture, then fixed with 4% PFA and stained with 1:5000 rabbit anti-EGFP (Origene) and anti-rabbit AF647 (10*μ*g/ml) following culture. Coverslips were mounted with mounting medium containing DAPI (Vector) on glass slides and imaged on a Nikon A1 Confocal microscope fitted with a 20x, 40x, or 60x objective.

### Western blot

For protein analysis, placental tissues or size exclusion column fractions were stored at −80°C until downstream analysis. Tissues were homogenized in RIPA buffer supplemented with 2 *μ*g/ml aprotinin, 5 *μ*g/ml leupeptin, and 1 mM phenylmethylsulfonyl fluoride (PMSF). Protein was quantified using a Qubit 3 fluorometer (Life Technologies, Carlsbad, MA). Samples were heat-denatured at 98°C for 15 minutes in RIPA buffer containing dithiothreitol (DTT). Denatured samples (10 *μ*g/lane) were run on precast 4-20% SDS-PAGE gels (Bio-Rad, Hercules, CA) and protein was transferred onto PVDF or nitrocellulose membranes. Membranes were blocked for 20-30 minutes in SuperBlock solution (ThermoFisher, Rockland, IL) at room temperature before overnight incubation at 4°C in 3% skim/tris buffered saline (TBS) milk with anti TSG101, anti-ITG alpha V (Abcam), anti-CD9 (Thermo Fisher), anti-ITG alpha 3, anti-ITG alpha 5, anti-ITG alpha 6, anti-ITG beta 1, anti-ITG beta 3 (Cell Signal Technologies), or anti EGFP (Origene) primary antibody. Membranes were washed three times for 5 minutes in TBS with 0.05% Tween-20 (TBS-T) at RT and incubated with 1:2000 horseradish peroxidase (HRP) conjugated goat anti-rabbit secondary antibody for 1hr at RT. Membranes were washed three times with TBS-T for 5 min and incubated with SuperSignal PicoPlus chemiluminescent substrate for 5 min before being imaged on a digital iBright membrane imager.

### Fluorescence/immunofluorescence microscopy and EV quantification in tissues

Tissues were harvested from mice were fixed in 4% paraformaldehyde (PFA) [Sigma-Aldrich, St. Louis, MO] in PBS and 30% sucrose in PBS at 4°C overnight before being frozen with isopentane and liquid nitrogen in Tissue Tek O.C.T embedding compound (Sakura Finetek, Torrance, CA). For fluorescence microscopy, 5 *μ*m tissue cryosections were mounted with DAPI nuclear stain and imaged on a Nikon Eclipse Ti epifluorescent microscope fitted with a 20x NA 0.5 plan fluor objective using Nikon NIS-Elements AR 4.40 software. To quantify EV in tissues, five random fields were imaged, and individual color channels processed using ImageJ. PKH26 and nuclei counts were computationally quantified using a custom pipeline developed in CellProfiler 3.0 software. For immunofluorescence microscopy, slides were stained with BioLegend antibodies and mounted with mounting medium with DAPI.

### Tissue clearing and immunofluorescence analysis

For clearing tissues prior to 3D imaging, paraformaldehyde-fixed organs were embedded in 2% agarose. Sections (200*μ*m) were obtained using a Leica VT1200 S vibratome (Leica, Buffalo Grove, IL), placed in 12-well plates, and permeabilized in 0.05% tween 20. Sections were blocked in 10% SuperBlock solution for 4-12 hrs, and stained with antibodies at 37°C in an orbital shaker for 24 hours. DAPI (10ug/ml) was added to the solution and sections were incubated for an additional 24 hours. After a 12-hour wash, secondary antibody was added overnight at 37°C, and samples were again washed for 12 hrs. Samples were then immersed in 2-3ml of CUBIC1 solution and incubated overnight at 37° before being washed briefly with PBS before immersed with 2-3ml of CUBIC2 solution for 12hrs.^24^

For traditional immunofluorescence, paraformaldehyde-fixed tissues were placed in 30% (W/V) sucrose solution in PBS overnight before being embedded in optimal cutting tissue (O.C.T.) compound (Sakura Finetek, Torrance, CA) and freezing. Cryosections (7*μ*m) were placed onto charged glass slides, blocked with 10% SuperBlock and 3% goat serum, and incubated overnight at 4°C with rabbit anti-EGFP followed by goat anti-rabbit conjugated to Alexa fluor 647. Slides were washed three times with 0.05% Tween PBS (PBS-T) for 5 min, and sections were mounted in DAPI-containing medium (Vector, Burlingame, CA).

All tissues and cells were imaged on a Nikon A-1 confocal microscope fitted with a 20X and 40X oil objectives lenses and images were processed using FIJI image analysis software. Images of the 200*μ*m vibratome sections were also processed using the Imaris v9.2 software (Bitplane) for 3D reconstruction of vasculature and EV localization. 3D mesh of CD31 positive vasculature was created using the Surface plugin and background subtraction was applied. Placental EVs were detected using the Spots plugin based on PKH26 labeling. The diameter of EV was estimated to be 0.75um in the XY slice for creating spots. Images were taken using the Snapshots function and 3D Video was generated using the animation function of the software.

### Flow cytometry

Lungs were minced and incubated in digestion buffer consisting of RPMI 1640, 0.05% liberase TH (Roche), and 100U/ml DNase1 (Sigma-Aldrich, St. Louis, MO) for 1 hr at 37°C. Larger tissue pieces were further dispersed by passing through a 3 ml syringe with a 27 gauge needle and cells were passed through a 70 micron nylon cell strainer (Falcon, Tewksbury, MA). The enzymatic digestion was quenched with 5% bovine serum albumin in PBS and cells were pelleted at 500 x g for 5 minutes before treatment with ACK lysis buffer (150mM NH4, 10mM KHCO3, 0.1mM Na2EDTA, in distilled water) to lyse red blood cells. Cells were resuspended in FSB (2% fetal bovine serum and 0.01% sodium azide in PBS), and staining was performed in U-bottom 96 well plates. Cells were incubated with Live/Dead Yellow (1:1000) followed by Fc block (1:200; BioLegend, San Diego, CA), and stained with antibody cocktails for 30 minutes on ice. Cells were washed twice and fixed with 1% PFA in flow staining buffer. Spectral flow cytometry was performed on an Aurora (Cytek BioSciences, Fremont, CA) with eUltracomp compensation beads used for single color controls. Events (100,000-200,000) were collected for each well and analysis was done using Kaluza 2.1 software (Beckman Coulter, Brea, CA). Flow cytometry t-distributed stochastic neighbor embedding (t-SNE) analysis was performed using the Cytofkit R package^56^ with default settings.

To characterize BMDC expression of mTomato or mGFP fluorescence, co-cultured dendritic cells were removed from culture plates with cell scrapers and stained with live/dead yellow viability stain (L34959, Thermo Fisher, Waltham, MA) for 15 min on ice. Cells were then blocked with TruStain FcX block (Biolegend, San Diego, CA) for 10 min and stained with anti-CD45, CD11c, and MHC-II antibodies (BioLegend, San Diego, CA) as described above. Single color controls were made with individual antibodies along with eUltracomp beads (Thermo Fisher, Waltham, MA), and spleen cells from mTomato and mGFP mice were used as endogenous fluorescence controls. Stained cells were analyzed by an Aurora spectral flow cytometer (CytekBio, Fremont, CA), and data were analyzed by Kaluza flow cytometry software (Beckman Coulter, Pasadena, CA).

### Whole organ imaging

To quantify EV localization in whole organs, 2.5 x 10^10^ GD14.5 DiR labeled pEVs were injected intravenously into NP female mice. After 24 hours, mice were euthanized and the lung, liver, spleen, thymus, brain, uterus, spleen, lymph node, kidney, and heart were harvested and fixed in 4% PFA in PBS for four hours before being stored in PBS. Whole organs were imaged on a LiCor odyssey infrared imager on manual scan mode (21 microns) under automatic acquisition settings (LiCor BioSciences, Lincoln, NE). Raw tifs were analyzed using Fiji and converted to 8 bit before mean intensity was measured in each tissue.

### Recombination genotyping

To detect genomic recombination in co-cultured mT/mG dendritic cells, DNA was isolated with a Quick DNA/RNA miniprep kit (D7005, Zymo, Irvine, CA) following the manufacturer’s instructions. DNA was amplified using a modified two-step polymerase chain reaction protocol targeting the unrecombined mTomato locus or recombined mGFP locus in mT/mG reporter (Figure 7C) mice.^31^ The mTomato locus was amplified with mT-F 5’-GCAACGTGCTGGTTATTGTG-3’ and mT-R 5’-TGATGACCTCCTCTCCCTTG-3’ primers, yielding a 200bp amplicon. The mGFP locus was amplified with mG-F 5’-GTTCGGCTTCTGGCGTGT-3’ and mG-R5’-TGCTCACGGATCCTACCTTC-3’ primers, yielding a 376 amplicon. Genomic DNAfrom dendritic cells was amplified for 35 cycles, and 3*μ*l of the PCR product was used as the template for a second round of PCR. Genomic tail DNA from WT, mTomato, and mGFP mice were used as positive and negative controls for both PCR reactions. PCR products were run on 1.5% agarose gel with 100bp ladder (New England Biolabs, Cambridge, MA) and visualized on an iBright digital gel imager (Thermo Fisher, Waltham, MA).

### Statistical analysis

For boxplots, the middle line represents median value, upper and lower box regions correspond to third and first quartiles (75th and 25th percentiles) and whiskers represent 1.5 times the interquartile range. All plots and analyses were generated in R v4.0. All raw data, analysis, and scripts for generation of figures are available on GitHub. Fiji/imageJ macros and CellProfiler pipelines are available on GitHub. Data were subjected to a Shapiro normality test before selecting the appropriate parametric or non-parametric statistical test.

## ACKNOWLEGEMENTS

The authors thank lab members including Geoffrey Grzesiak, Sarika Kshirsagar, and Cole McCutcheon for technical assistance and useful discussions; Alicia Withrow for assistance with transmission electron microscopy; and Melinda Frame for imaging assistance in confocal microscopy.

## Key Resources Table

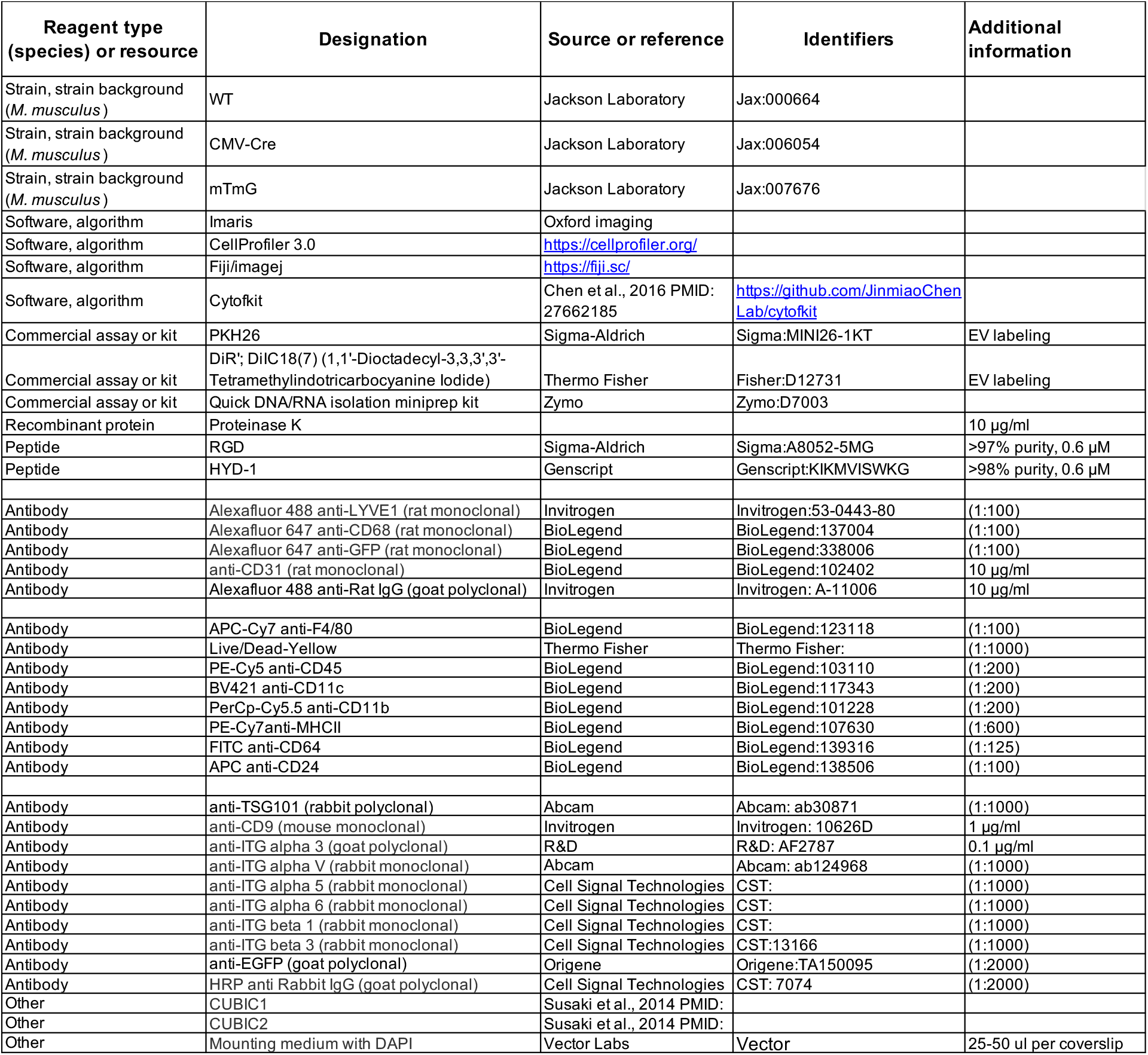

**Supplemental Figure 1.**
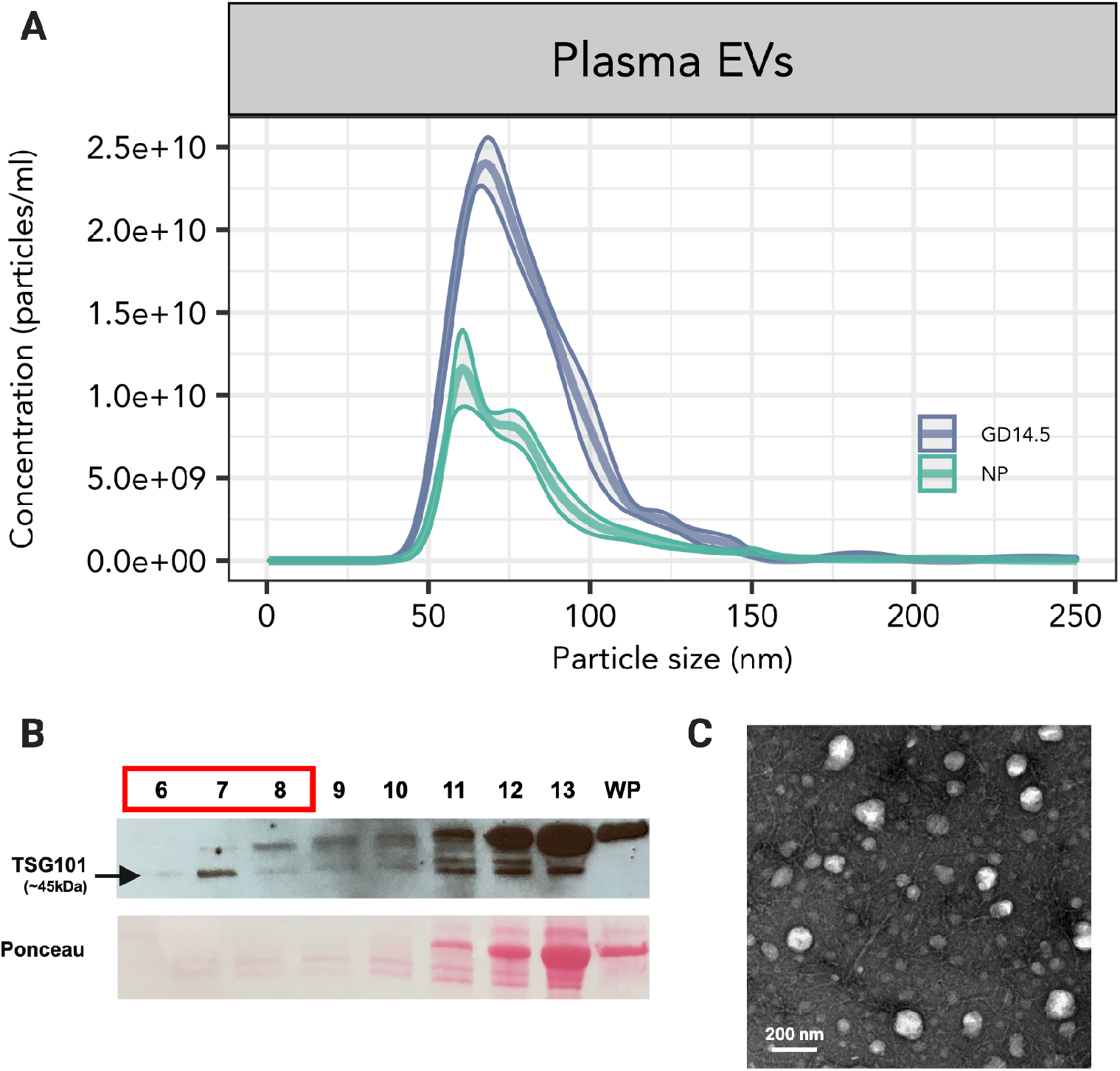
Isolation and validation of plasma EVs. A. Representative concentration and size distribution analysis of plasma EVs from non-pregnant (NP) and gestational day (GD) 14.5 mice obtained by ultracentrifugation and analyzed by NanoSight. The thick middle line in each distribution curve represents mean concentration, and the shaded area bound by narrower lines represents SEM. Data represent five technical replicates of three different animals in each group. B. Western blot analysis of plasma EVs from a GD14.5 mouse. Lane numbers represent size exclusion chromatography fractions; red box indicates fractions used for in vivo experiments. WP, whole plasma. Representative data from three independent experiments. C. Transmission electron microscopy of plasma EVs (fractions 6-8) from a nonpregnant mouse (representative image of n = 3).

**Supplemental Figure 2.**
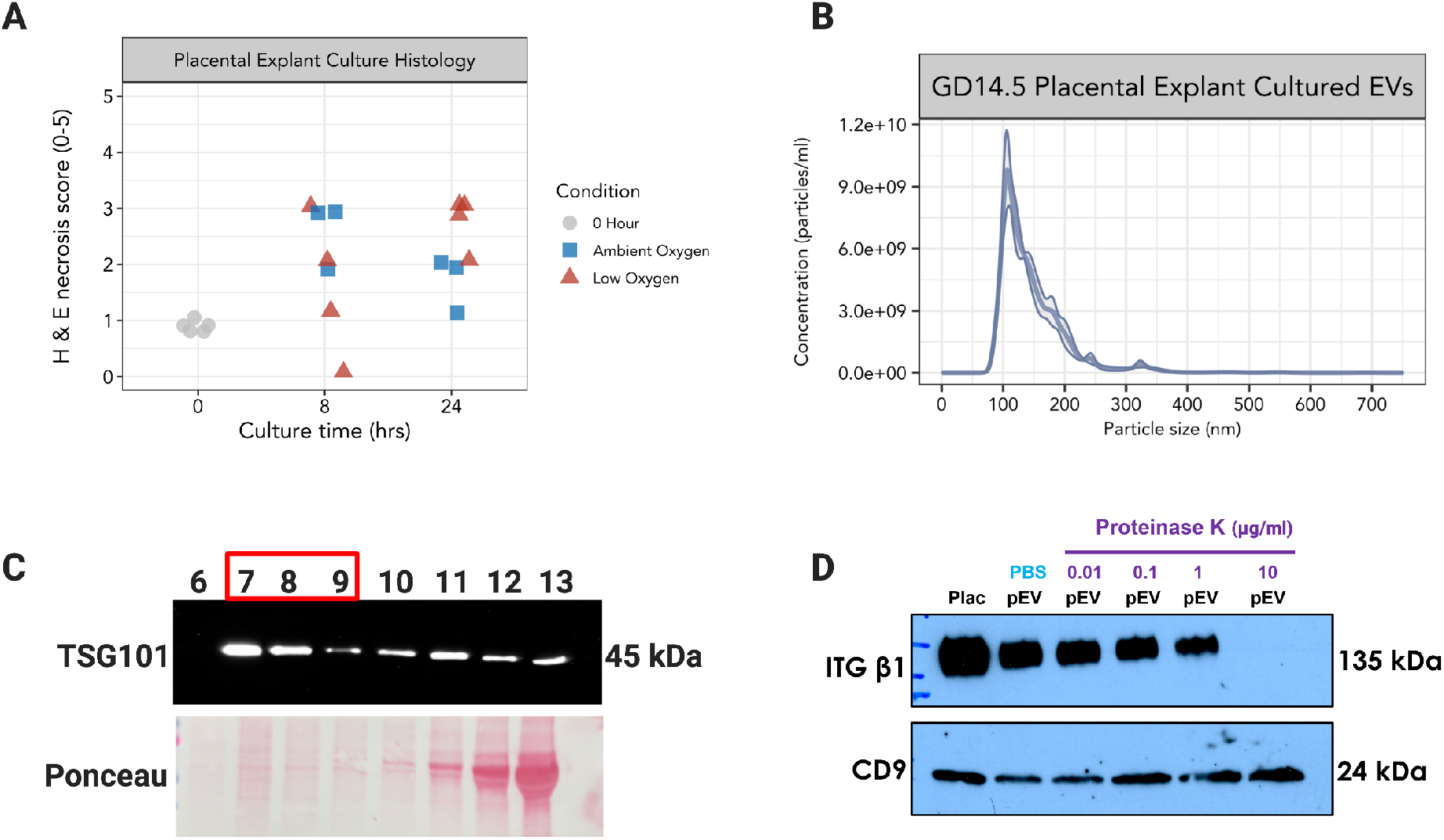
Validation of placental explant culture. A. GD14.5 placentas were from GD14 were cultured under ambient or low (8% oxygen) conditions for 8 or 24 hours and processed for histological analysis. Uncultured placentas were dissected and processed immediately after dissection (0 hr). Hematoxylin and eosin-stained sections were subjected to blinded scoring for degree of necrosis on a scale of 0-5. B. Representative histogram of placental EVs analyzed by nanoparticle tracking analysis. C. Western blot analysis of placental EVs from culture supernatants. Numbers over lanes represent fractions from size exclusion chromatography; red box indicates fractions used for in vivo experiments. D. Western blot of placenta (plac) and placental EVs treated with PBS (negative control) or increasing concentrations of proteinase K. ITG*β*1, which was strongly expressed by placental EVs, was efficiently removed by 10 μg/ml proteinase K, which did not remove immunoreactivity for the exosomal marker CD9 (SFig. 2D). Blot is representative of three independent experiments.

**Supplemental Figure 3.**
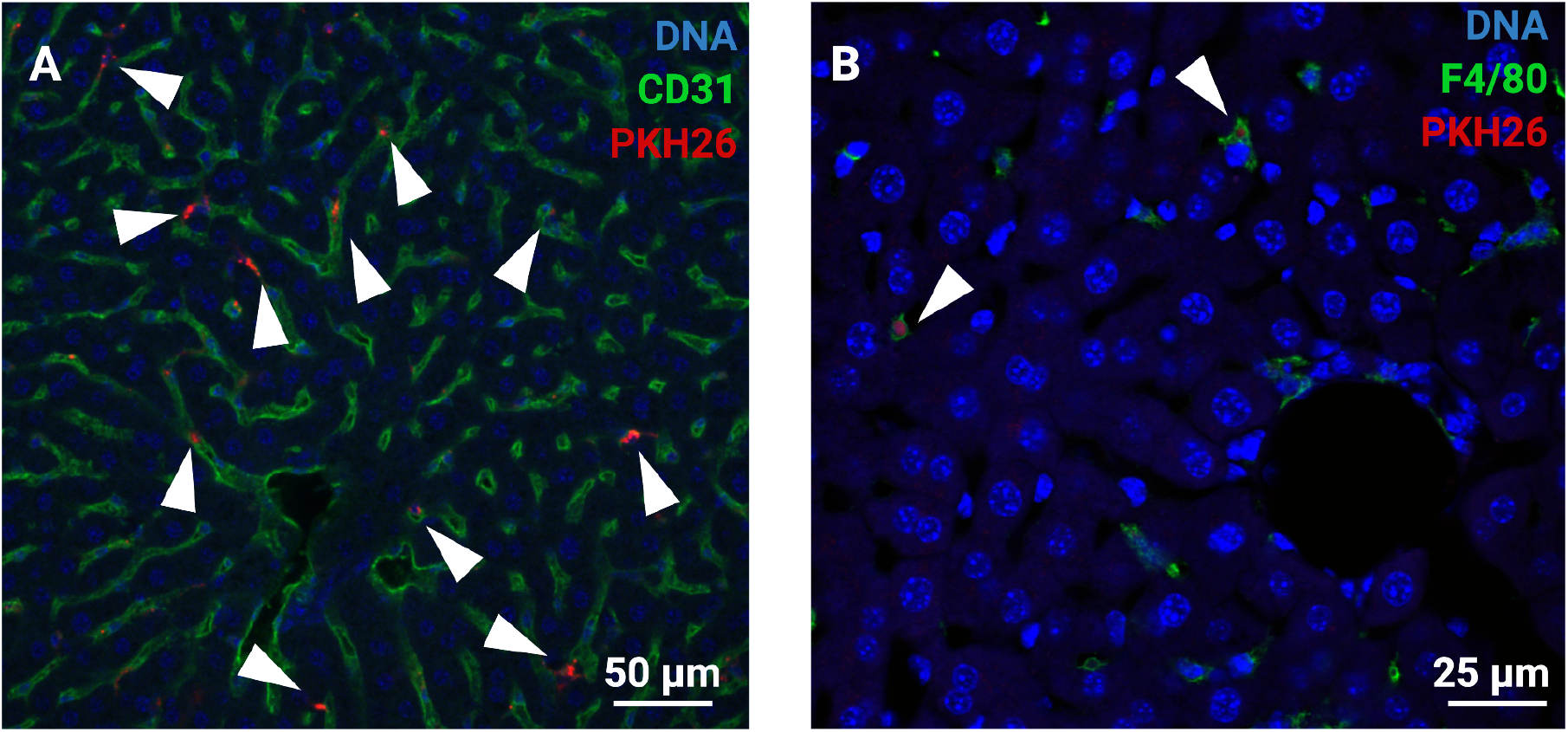
Placental EV Localization in Liver. A. Immunofluorescence confocal microscopy showing colocalization of placental EV (red punctate fluorescence) with CD31+ endothelial cells. B. Immunofluorescence colocalization of placental EV with F4/80+ macrophages Arrowheads highlight colocalization. Representative images from three independent experiments.

**Supplemental Figure 4.**
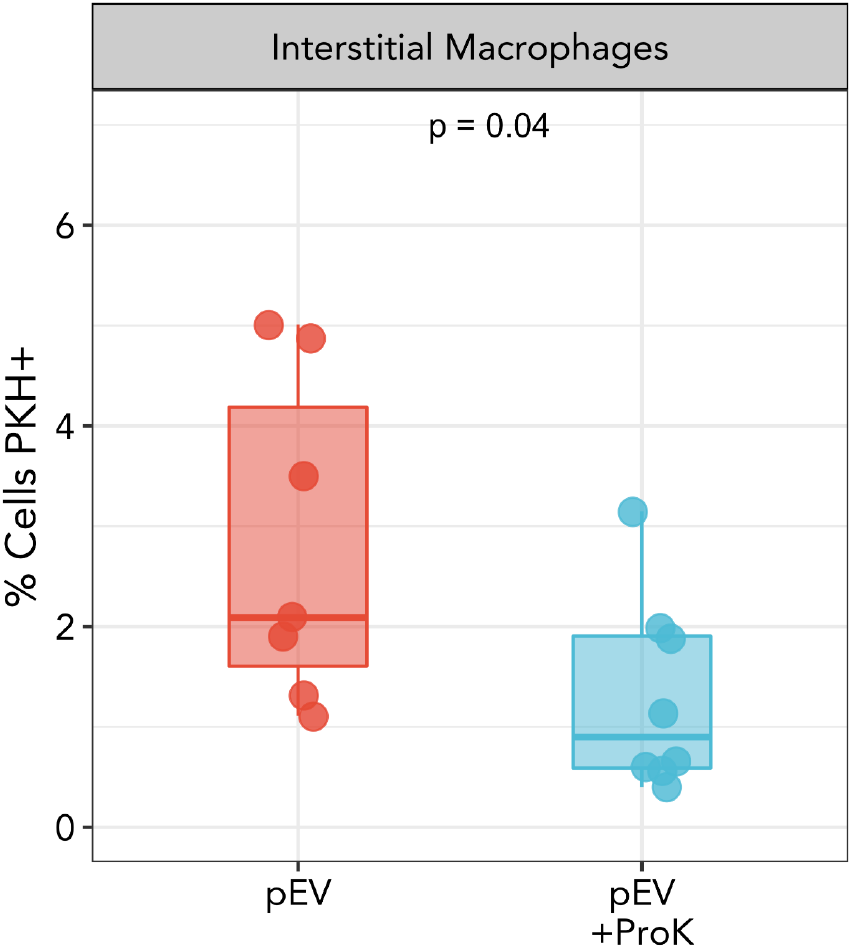
Proteinase K inhibits pEV localization to lung interstitial macrophages. Flow cytometric quantification of placental EV localization to interstitial macrophages in lung. Points represent individual biological replicates; data were analyzed by Welch’s T-test.

**Supplemental Figure 5.**
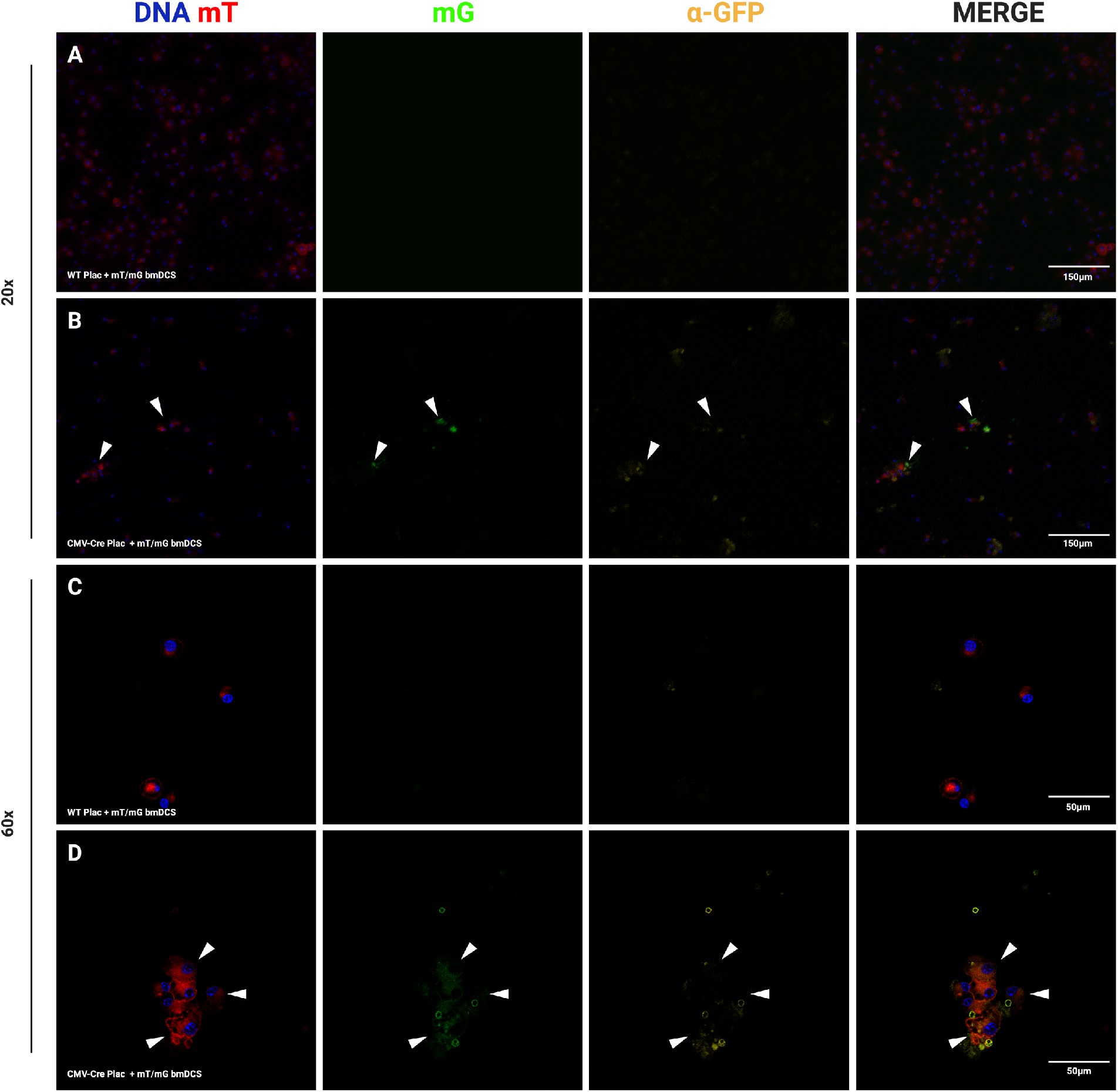
In vitro recombination of BMDC co-cultured in the presence of CMV-Cre placentas. Reporter BMDC from mT/mG mice were co-cultured with WT or CMV placentas and viewed by confocal microscopy for absence (mT) or presence (mG) of recombination. Results of mG were confirmed using an AF-647-conjugated anti-GFP antibody (a-GFP). Representative confocal microscopy of reporter BMDCs co-cultured with WT (left) or CMV-Cre (right) placentas. Arrowheads point to GFP foci in cells. A, C: BMDC cocultured with WT placentas. B, D: BMDC cocultured with CMV-Cre placentas.

